# pH-Responsive Phase Separation Dynamics of Intrinsically Disordered Peptides

**DOI:** 10.1101/2025.01.09.632076

**Authors:** Manali Nandy, Ketan A. Ganar, Hans Ippel, Ingrid Dijkgraaf, Siddharth Deshpande

## Abstract

Liquid-liquid phase separation of biomolecules is crucial for maintaining the functional organization in biological systems. Intrinsically disordered proteins are particularly prone to form phase-separated condensates in response to various physicochemical triggers. While the effect of ionic strength and temperature on phase separation dynamics have been studied extensively, the influence of pH is less explored. Here, we study a model glycine-rich protein present in the tick bioadhesive, given its capability to undergo phase separation. After confirming its disordered nature through spectroscopy, we investigated its pH dependence and underlying molecular mechanisms. Our findings reveal that pH significantly influences the protein hydrophobicity via ionic residues, driving notable variations in the coacervation behavior (propensity, progression) and in shaping the material properties (viscosity, interfacial activity) of the formed condensates. Given the ubiquitous presence of disordered proteins in biology, this study provides valuable insights about the broad implications of the pH-dependent behavior of intrinsically disordered proteins.

## Introduction

Liquid-liquid phase separation (LLPS) of biomolecules is increasingly recognized as a fundamental process in living systems^1,2^. Associative LLPS is manifested in the form of biomolecular condensates^3^, which arises via coacervation, where a biopolymer-rich condensed phase co-exists in dynamic equilibrium with the biopolymer-poor dilute phase. Beyond their role in cellular function and organization, including those of membraneless organelles^4–6^, condensates have diverse applications such as drug delivery carriers^7^, tissue culture scaffolds^8^, novel adhesive materials^9,10^, and developing synthetic cells^11^. The formation of biomolecular condensates is dependent on the multivalency as well as the flexibility of biopolymers such as proteins and nucleic acids^12,13^. Intrinsically disordered proteins/peptides (IDPs) are important classes of biomolecules which structurally lack independently folded domains, making them prone to LLPS^14–16^. These flexible IDPs act as random coils, allowing the functional groups of amino acids to easily form weak multiple bonds with neighboring groups^13,17–19^, acting as the primary driving force for LLPS. For instance, the electrostatic interaction between acidic and basic amino acids^20,21^, hydrogen bonding^22^, hydrophobic interactions^23^, cation-π interactions between the basic and aromatic amino acids^24^, and π- π interactions among the aromatic amino acids^25,26^ contribute to the condensation of IDPs. The rapid association and dissociation of multiple intermolecular and intramolecular bonds between IDPs is responsible for the fluid-like nature of condensates^27,28^. Furthermore, the material properties of condensates can change over time leading to the formation of more solid-like structures^29,30^. Over the years, extensive research has focused on now well-known IDPs such as tau^31,32^, DDX4^2,33^, hnRNPA2^34,35^ , FUS^36,37^, CAPRIN1^38,39^ which undergo LLPS under various biological triggers.

Recently, we explored the phase separation potential of a tick salivary protein belonging to the glycine-rich protein (GRP) family^40^. GRPs are an important class of IDPs^41,42^ and have kept their footprints throughout several biological systems, from intracellular organelles^43,44^ to extracellular secretions^45,46^. Tick saliva is rich in GRPs^47–50^ and undergoes liquid-to-solid transition to form a strong bioadhesive, also known as the cement cone, ensuring the tick’s firm attachment to its host. We showed that a tick GRP, which we termed tick-GRP77 (Uniprot ID: Q4PME3), undergoes coacervation in presence of kosmotropic salts and further experiences liquid-gel transition over time. Our findings revealed that cation-π and π-π interactions between the aromatic and arginine amino acid residues play crucial role in the LLPS of tick-GRP77 and that the formed condensates are highly adhesive in nature. Encouraged by the observed salt-dependent tick- GRP77 phase transitions, we considered other physicochemical changes that the tick saliva encounters during the cement cone formation and identified pH as an important parameter. While the tick saliva is quite basic (pH 9.5)^51^, it encounters a relatively acidic host environment (skin pH ≈ 5^52,53^; blood pH ≈ 7.4^54^). Furthermore, while extensive research has been carried out on the effect of salt^55,56^, temperature^57,58^, and crowding agents^59,60^ on LLPS of several IDPs, the impact of pH on condensates remains underexplored. As a result, fundamental understanding of the material properties and thorough structural analysis regarding the role of pH on intermolecular interactions between the amino acids is still scarce.

In this work, we present a rich and nuanced, pH-dependent coacervation behaviour of intrinsically disordered peptides, with tick-GRP77 serving as a model system. We first confirmed the highly disordered nature of tick-GRP77 using nuclear magnetic resonance (NMR) spectroscopy. We then systematically investigated the coacervation dynamics in response to a broad pH range using microscopic observations and various physical chemistry tools, including droplet evaporation assay, fluorescence microscopy, and tensiometry measurements. Our findings reveal that the degree of hydrophobicity of IDPs varies significantly with changing pH and directly manifests into coacervation dynamics and the material properties of the resulting condensates. This study thus provides valuable insights into the molecular mechanisms underlying pH-dependent phase separation and highlights the broader implications for the behavior of intrinsically disordered proteins in diverse biological contexts.

## Results

### Sequence and spectroscopic analyses show that tick-GRP77 is pH-sensitive and highly disordered

The mature tick-GRP77 amino acid sequence, omitting the signal peptide^40^, is shown in **Figure 1a**. As can be seen, the sequence is rich in glycine residues (G), and is interspersed with cationic (lysine – K, arginine – R, and one histidine – H) and aromatic (tyrosine – Y and phenylalanine – F) residues. It is worth noting that owing to its imidazole group, H is capable of exhibiting both cationic and aromatic properties^61^. The periodic positioning of the aromatic and cationic residues resembles the spacer-sticker model which facilitates cation-π and π-π interactions leading to LLPS^62,63^. Furthermore, the N-terminus, being abundant in ionic amino acids (K, glutamic acid – E, and aspartic acid – D), is more hydrophilic than the C-terminus that is rich in cationic (R, H) and aromatic (F, Y) residues, responsible for cation-π and π-π interactions^40^. We synthesized tick-GRP77, and its two separate halves (N- and C-terminus, as shown in **Figure 1a**) via solid-phase peptide synthesis and native chemical ligation (see Methods for details). A schematic representation of tick-GRP77 undergoing LLPS owing to the cation-π and π-π interactions and a typical microscopic image of the formed condensates in presence of phosphate buffer saline (PBS, pH 7.4) during a droplet evaporation assay (see Methods for details; also see the next section) is shown in **Figure 1b**. Fluorescently labelled tick-GRP77 (OG488-GRP77; fluorescent label: Oregon Green 488; 5 mol%) was added for visualization.

**Figure 1:**
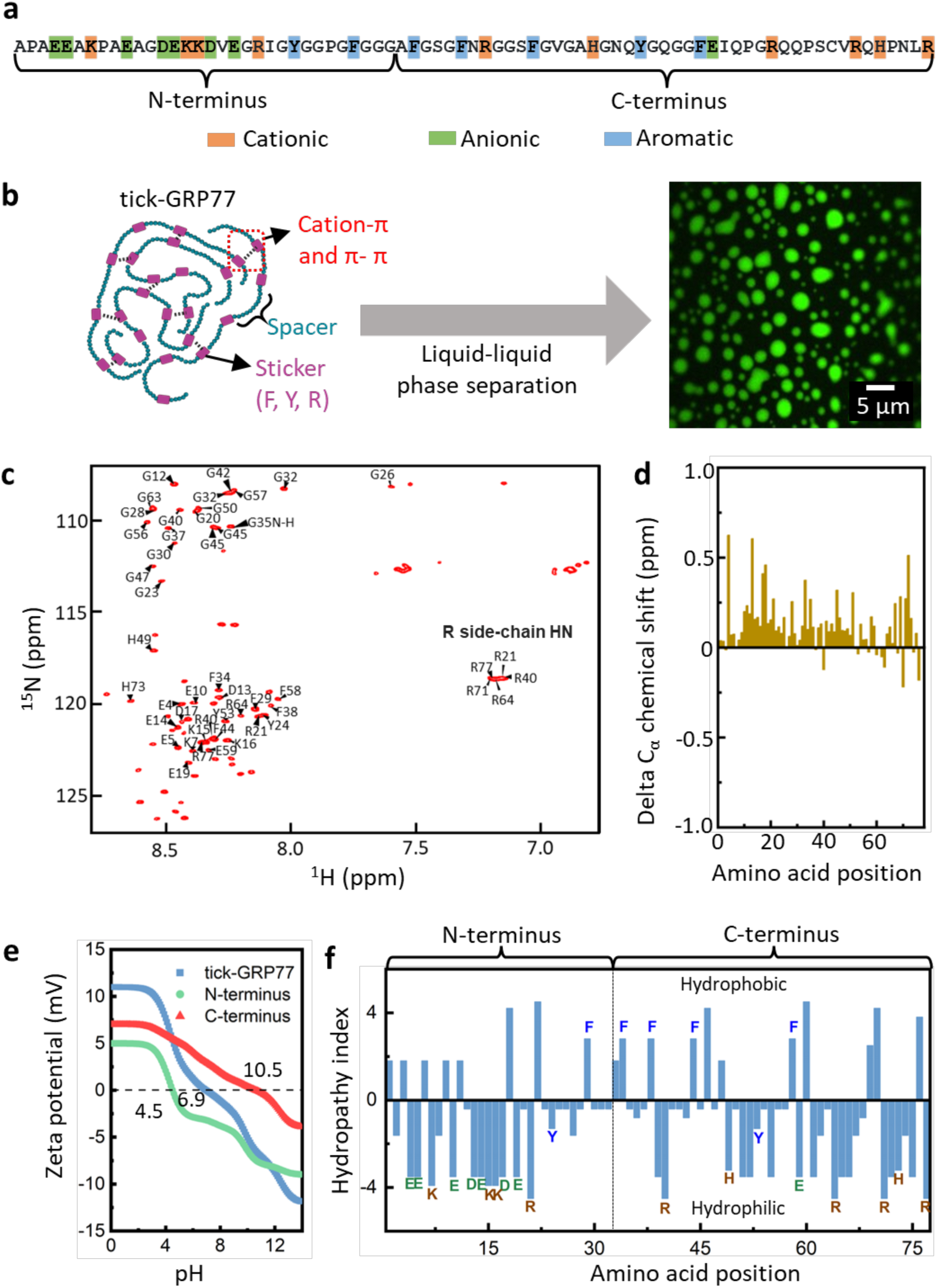
tick-GRP77 is intrinsically disordered and displays pH responsiveness. (**a**) Amino acid composition of tick-GRP77, showing its glycine-rich nature; pH-sensitive cationic and anionic residues, and aromatic residues responsible for phase separation are highlighted. The two halves, N- and C-terminus, are also indicated. (**b**) Schematic overview of tick-GRP77 undergoing phase separation owing to its sticker- spacer structure and cation-π and π-π interactions. tick-GRP77 condensates formed by droplet evaporation in presence of PBS, with OG488-GRP77 (5 mol%) added for fluorescence visualization. (**c**) Natural abundance 700 MHz 2D ^15^N-^1^H HSQC spectrum of tick-GRP77 recorded in water at low pH (3.8) and at 16 °C. Amide cross peaks in the spectrum belonging to individual amino acid have been assigned. Amide peaks instrumental for coacervation (G, R, K, F, Y), plus those amenable to pH response (D, E, K, R, H), are labelled by residue sequence number. (**d**) Experimentally derived delta C_α_ chemical shift values for tick-GRP77 relative to predicted random coil shifts show low values near zero ppm, indicating the random coil nature of the peptide. (**e**) Theoretical titration curves showing the distinct pI values of tick-GRP77 and its two halves. (**f**) Hydropathy index of tick-GRP77 based on Kyte-Doolittle hydropathy model with the cationic, anionic, and aromatic amino acids clearly marked. The N-terminus is rich in ionic residues making it more hydrophilic than the C-terminus which is rich in aromatic and cationic residues.

The intrinsically disordered structure of tick-GRP77 is already predicted by several disorder prediction algorithms^40^. We experimentally confirmed this using NMR spectroscopy (**Figure 1c-d**, see Methods for details). We particularly looked at the NOE (nuclear Overhauser effect) contacts in the 2D NOESY (nuclear Overhauser effect spectroscopy) NMR spectrum. In the 2D NOESY NMR spectrum, only short-range NOE peaks were observed with no long-range NOE peaks (**Figure 1c**), which is consistent with a mostly unstructured random coil-like protein conformation^64^. The NOE peaks corresponding to the pH-sensitive amino acids (D, E, R, K, H) and the ones crucial to condensation (R, K, F, Y, and G) are marked along with their position numbers. This was further corroborated with the analysis of experimentally determined C_α_, C_β_, H_α_, ^15^N, and HN chemical shifts (𝛿, in ppm) over the entire sequence range of the protein utilizing the POTENCI random coil prediction tool^65^. These POTENCI-generated simulated chemical shifts are derived from a large set of experimental model peptides in which predicted sequence-dependent random coil chemical shift index (CSI) values are recalculated as a function of pH and temperature. **Figure 1d** displays the delta C_α_ chemical shifts (𝛿_𝑒𝑥𝑝𝑒𝑟𝑖𝑚𝑒𝑛𝑡𝑎𝑙_ − 𝛿_𝑟𝑎𝑛𝑑𝑜𝑚 𝑐𝑜𝑖𝑙_) at pH 3.8 between experimental values and POTENCI-predicted sequence-dependent values for tick-GRP77 against residue number. **Supplementary**

**Figure 1** show the C_β_, H_α_, ^15^N and HN delta chemical shift values. At pH 7, most amide resonances tend to be broadened by increased solvent exchange rate, but aliphatic resonance positions in the 2D spectra remain relatively constant as a function of pH, demonstrating similar structure at physiological pH. Moreover, elevating the solution pH of the NMR sample of tick-GRP77 to physiological pH (7–7.4) did not induce any secondary structure. The close agreement between predicted and experimental ppm values at pH 3.8 verifies the low complexity and absence of secondary structures in tick-GRP77. By averaging the delta C_α_ chemical shift values over the whole sequence, we estimated that at most 2.1% of the protein transiently adopts an 𝛼-helix state when time-averaged under low pH NMR conditions.

With the disordered nature of tick-GRP77 established, we proceeded with determining an appropriate pH range for studying the effect of pH on the phase separation behavior. Based on the clear amino acid variations for the two termini, we expected distinct pI values for the three peptides. We theoretically calculated the titration curves^66^, and as can be seen in **Figure 1e**, indeed obtained significantly different values of the isoelectric points (pI) for the full-length tick-GRP77 (6.9), the N - (4.5), and the C-terminus (10.5). This provided us with an appropriate system to systematically study the effects of pH across a broad yet physiologically relevant range of pH 4 to 10. We thus chose three distinct pH values: acidic (pH 4), neutral (pH 7), and basic (pH 10 or 12) to cover the entire pI range. For the C-terminus, we considered pH 12 instead of pH 10 as its pI is at 10.5.

As tick-GRP77 consists of several hydrophobic and hydrophilic amino acids, we used the Kyte and Doolittle hydropathy scale^67^ to analyze the hydrophilicity/hydrophobicity of the amino acids present in the sequence. As shown in **Figure 1f**, positive values portray hydrophobic nature, while negative values reflect hydrophilic nature of the individual residues. We note that the ratio of hydrophobic-to-hydrophilic amino acids for the three peptides is similar (close to 0.3), reflecting their intrinsic hydrophilic nature, making them apt for studying pH-responsiveness and its possible manifestation into coacervation dynamics and condensate properties.

### Droplet evaporation assay reveals the pH-dependent traits of IDP condensates

To assess the effect of pH on the phase separation of tick-GRP77 and its two separate termini, droplet evaporation assays (schematically shown in **Figure 2a**) of peptide solutions prepared in PBS were performed (see Methods for details). For all the droplet evaporation assays, we used a peptide concentration of 50 uM and selected three distinct pH values (4, 7, and 10/12) as previously described. A pH of 12 was specifically applied for the C-terminus due to its higher pI value. The pH was adjusted by the addition of HCl or NaOH; we opted to let go of the buffering capacity of PBS and did not use varying buffers for different pH values in order to keep the variability of the buffer components to a minimum. To begin the assay, we casted a 2 µL peptide solution on a microscopic glass slide and let it evaporate at room temperature. Since the evaporation flux is highest at the edge of the droplet, the peptides reached a critical concentration at the droplet boundary, leading to simple coacervation^68,69^. The rightmost panel of **Figure 2a** shows a two-dimensional representation of the droplet edge, where the LLPS process is manifested in two ways. First, a sharp phase boundary forms in the vicinity of the droplet boundary, resulting in a band of the condensate phase, which we refer to as ‘the rim’. Second, condensates develop in the droplet interior, that continuously fuse with the rim. We analyzed three key parameters emerging from the evaporation assay (see Methods for details): time required for the onset of coacervation (*t_i_*), i.e., the time interval between droplet casting and the formation of the phase boundary, rim thickness (*R_d_*), and the average diameter of the condensates (*d*).

**Figure 2:**
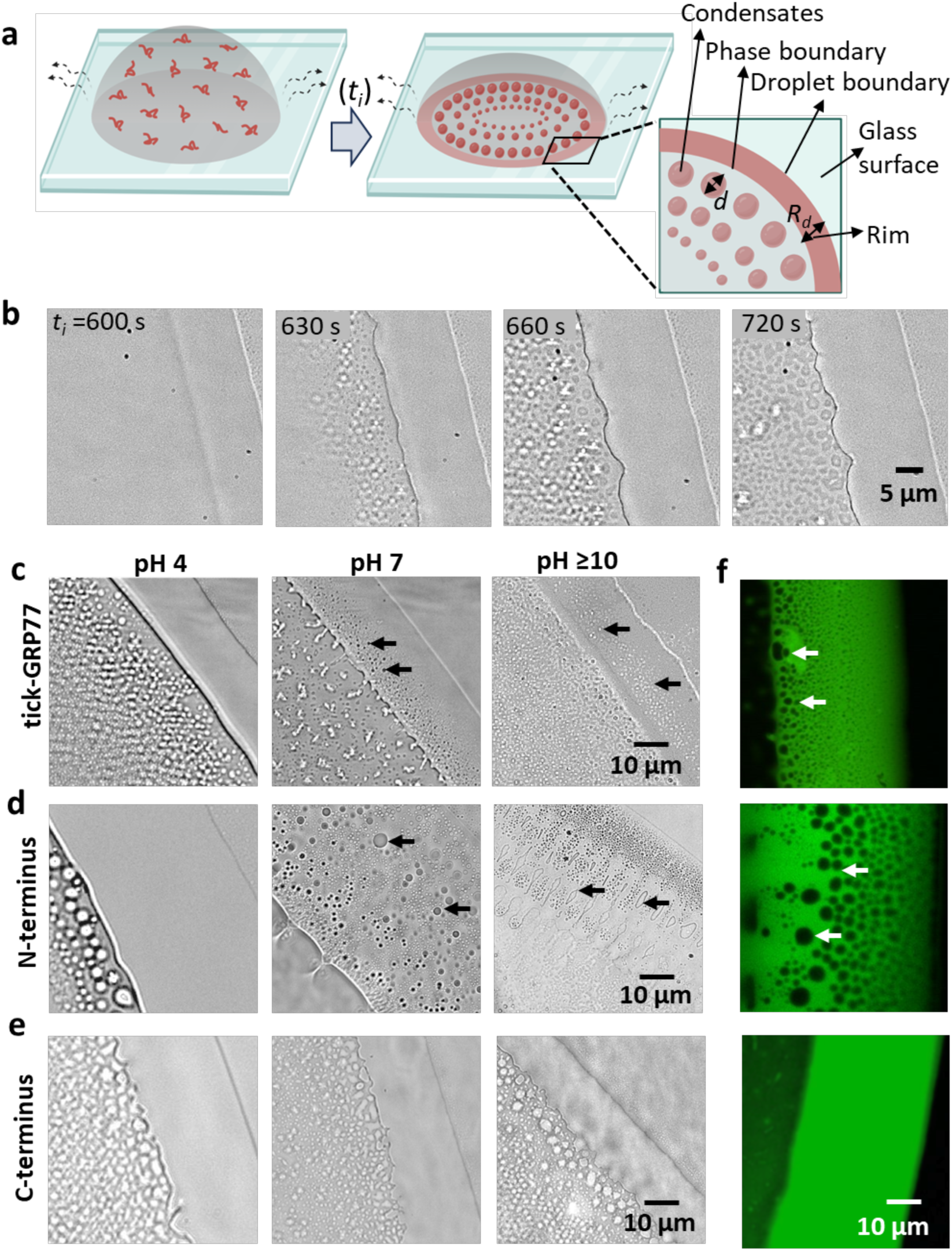
Droplet evaporation assay enabled to visualize the pH-based manifestation of the condensation behavior of tick-GRP77 and its two halves. (**a**) Schematic representation of a droplet evaporation assay, where a 2 µL-droplet consisting of 50 µM peptide in PBS is evaporated at room temperature. A close-up representation of the droplet edge showing the condensates, phase boundary, droplet boundary, and the protein-rich rim that is formed in between. The key parameters analyzed are highlighted: condensate diameter (d), rim thickness (R_d_), and onset of coacervation (t_i_). (**b**) A representative bright-field time-lapse showing the LLPS of tick-GRP77 at pH 4, manifested in the form a dense phase rim and condensate droplets forming in the interior and fusing with the rim. (**c**)-(**e**) Representative bright-field images of the droplet evaporation assay for tick-GRP77, N-terminus, and C-terminus, at pH 4, 7, and ≥ 10. Images were captured just prior to crystallization of salts in the sample. (**f**) Corresponding confocal images at pH 7. OG488-GRP77 (5 mol%) was added for fluorescence visualization. Arrows in (**c**), (**d**), and (**f**) indicate the protein-depleted aqueous regions inside the rim.

**Figure 2b** shows the time-lapse images of a typical droplet evaporation assay, taking the example of tick-GRP77 at pH 4. After approximately 10 min, a distinct rim appears and marks the onset of coacervation. Subsequently, the evaporation flux gradually becomes prominent within the droplet interior as well, and peptide condensates are generated in the central part of the droplet. Due to capillary flow, these droplets are transported towards the edge of the droplet^70^, where they tend to fuse with the rim. The rim thickness gradually increases, partially due to the fusion with the condensates. **Figure 2c** (and **Supplementary Videos 1, 2 and 3**) shows bright-field microscopy images of a small section of the droplet boundary for 50 µM tick-GRP77 at pH 4, 7, and 10 where the presence of rim and condensates can be seen simultaneously without pronounced differences in response to pH variation. We observed a similar response for the N-terminus as shown in **Figure 2d** (also see **Supplementary Videos 4, 5 and 6**) but with a much thicker rim. Interestingly, for tick-GRP77 and the N-terminus (at pH ≥ 7), we observed the formation of inverted phases within the rim, which are protein-depleted aqueous phases within the dense phase (marked by arrows in **Figures 2c-d**). These inverted phases were completely absent in case of pH 4. We, and others, have previously observed the presence of inverted phases within the rim during droplet evaporation assays^40,71^. We speculate that the origin of the inverted phase may arise from several factors: Firstly, the rim readily spreads on the glass substrate, occasionally engulfing the dilute phase during the process (**Supplementary Videos 5 and 6**). To aid to this, the rate of rim spreading could be significantly higher than the rate at which the condensate phase is generated. This may be further supported by higher peptide hydrophilicity, enhancing the water retention capability of coacervates, leading to lower compactness of peptides within coacervates and increased swelling^72^. An alternative explanation could be the protein concentration deviating from the binodal and entering the metastable spinodal region of the phase diagram as pH increases^4^, as reported for multiphase condensates during temperature cycling^73^.

Compared to the full peptide and the N-terminus, we observed a completely different response from the C-terminus, as can be seen from the corresponding bright-field images of the droplet evaporation assay of the C-terminus at different pH values in **Figure 2e** and in **Supplementary Videos 7, 8 and 9**. Unlike the other two peptides, C-terminus did not show the protein-depleted inverted phase in the rim at any pH values. Also, the phase boundary of C-terminus was rather poorly defined in the final minutes before crystallization. We took a closer look at the rim (for pH 7 cases) using confocal fluorescence microscopy and by adding 5 mol% OG488-GRP77 in the starting peptide solutions (**Figure 2f**). For tick-GRP77 and its N-terminus, the inverted phases were clearly visible (dark aqueous circular areas distributed in protein-rich highly fluorescent rim, marked by arrows), while a homogenous rim was observed for the C-terminus. The inverted phase will be discussed further in the next sections. We also used the confocal images to estimate the extent of partitioning of the peptides in the condensate phase. By calibrating the fluorescence intensity as a function of OG488 concentration (**Supplementary Figure 2**, see Methods for details), we estimated the tick-GRP77 concentration in the condensates to be ∼1500 times higher than in the dilute phase.

### pH-dependent coacervation behaviour is dictated by the ionic residues present in IDPs

In addition to the dependence of the inverted phase on pH and the peptide itself, we also looked at the three quantifiable parameters outlined earlier. The size of the condensates (*d*) across various conditions remained relatively consistent, with a size range of 0.5-2 µm, showing no significant trend (**Supplementary Figure 3**). In contrast, we observed that the rim thickness (*R_d_*) and the onset of coacervation (*t_i_*) values differed significantly between different peptides. **Figure 3a** shows the *t_i_* as a function of pH for all the three peptides. As the pH increased, *t_i_* steadily increased (by ≈ 60%) for the N-terminus, while it decreased by a similar extent (≈ 50%) for the C-terminus. For tick-GRP77, *t_i_* increased more modestly (≈ 18%), placing it in between the two peptide halves. Considering that the peptide concentration at the droplet boundary increases over time, an increase in *t_i_* can be associated with a lowered coacervation tendency. Similarly, when we analyzed the variation in *R_d_* as a function of pH for all the three variants (**Figure 3b**), we found a dramatically high *R_d_* for the N-terminus compared to the C-terminus as well as the tick-GRP77 across the entire pH range. Furthermore, the *R_d_* for the N-terminus increased two-fold while that for the C-terminus decreased two-fold, across the pH range. For tick-GRP77, a mild increase (1.5 fold) was observed, again reflecting an intermediate behavior between the two termini.

**Figure 3:**
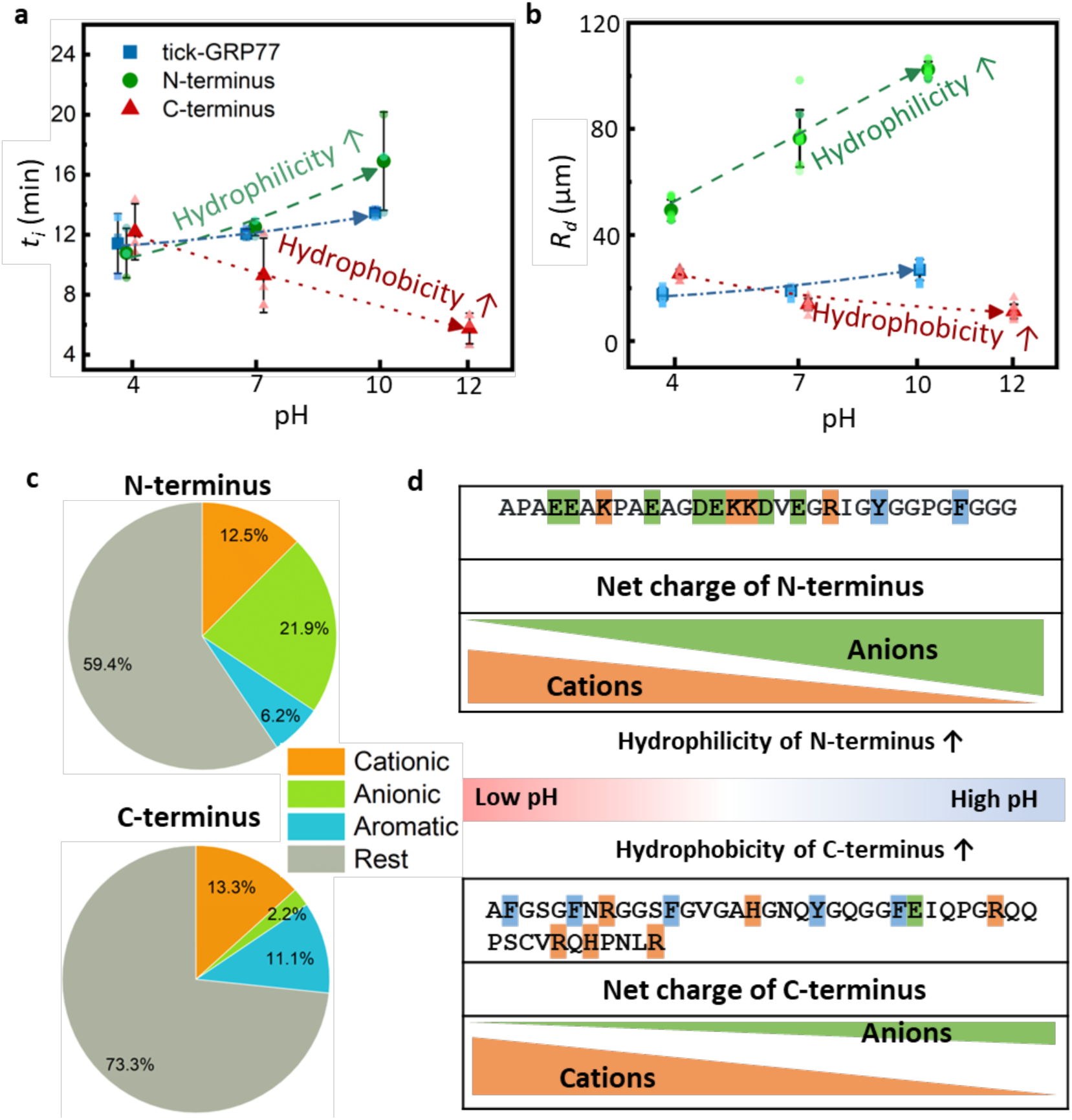
Onset of coacervation and rim thickness has its roots in the pH-responsive ionic residues present in IDPs. (**a**) Plot showing the onset of coacervation for tick-GRP77, C- and N-termini as a function of pH. The prominent increase and decrease in the t_i_ respectively for the N- and the C-termini hint to their corresponding increase in the hydrophilicity and hydrophobicity with pH increment. (**b**) Variation in the rim thickness for tick-GRP77, C- and N-termini as a function of pH. The increase and decrease in the R_d_ respectively for the N- and the C-termini again indicate their increasing hydrophilicity and hydrophobicity with pH increment. (**c**) Pie charts showing the percentage of cationic, anionic, and aromatic residues present in the two termini. N-terminus is rich in cationic as well as anionic residues. In contrast, C-terminus has higher content of aromatic residues, with very low fraction of anionic residues. (**d**) Schematic overview of the prominent changes in terms of the net charge for the two termini in response to pH variation and the overall trend in the hydrophilic/hydrophobic character of the peptide. The starting concentrations for all the droplet evaporation assays were 50 µM peptide in PBS. The plots in **a** and **b** are represented as mean ± s.d., along with individual data points (n = 3 independent experiments in each case). For improved visualization, the pH values are shifted by 0.2 pH units for the three IDPs, and the trends are indicated via dashed or dotted arrows.

Based on these observations, we wondered how the biochemical properties of the peptides could determine the observed phase separation behavior. Therefore, we took a more detailed look at their amino acid compositions, as shown in **Figure 3c**. It appeared that the N-terminus is abundant in anionic (21.9%) and cationic (12.5%) residues, with a relatively small fraction (6.2%) of aromatic residues, whereas the C-terminus is rich in cationic (13.3%) and aromatic (11.1%) residues and contains only one anionic residue (E). The analytical diagram in **Figure 3d** considers the charges at play in response to the pH variation. At very low pH, the charge of anionic residues become negligible owing to their low isoelectric points (pKa values of 3.65 for D and 4.25 for E^74^) and the cationic residues are fully charged due to their higher isoelectric points (pKa values of 10.53 for K, 12.48 for R, and 6.0 for H^74^). As the pH increases, the anionic side groups become more charged due to increasing deprotonoation of the COOH groups while the cationic side groups turn weaker owing to the gradual deprotonation of the NH_3_^+^ and imidazole groups. In case of the N-terminus, the proportion of π-electron-rich residues (F, Y, R, H) is low and they are involved in cation-π and π-π interactions during phase seapration. Hence, one can say that the N-terminus becomes more hydrophilic with increasing pH. In case of the C-terminus with negligible anion content, as the pH increases, the capacity of cationic residues participating in electrostatic interactions reduces. However, the cation-π and π-π interactions still hold strong, facilitated by more hydrophobic R residues instead of the K residues present in the C-terminus^62^. Additionally, the two H residues fully lose their cationic nature and contribute more towards hydrophobic interactions. Hence, the overall charged contribution from cations reduces and consequently, one can conclude that the C-terminus becomes more hydrophobic with increasing pH.

Thus, based on the contributions of present amino acid residues, we propose that as the pH increases, N-terminus is rendered more hydrophilic, while the C-terminus becomes more hydrophobic. As a matter of fact, the trends that we observe agree closely with this argument. To start with, more hydrophobic polymers, in principle, have a stronger tendency to coacervate because of the increased inter- and intramolecular interactions, thus explaining the decreasing *t_i_* for the C-terminus with increasing pH and the opposite in the case of N-terminus. Indeed, several reports have highlighted the higher propensity of coacervation for hydrophobic polymers compared to hydrophilic ones^75–77^ . Next, with the glass slide also being hydrophilic, the more hydrophilic N-terminus condensate phase tends to wet the glass slide and spreads rapidly, leading to a thicker rim. On the other hand, as the hydrophobicity increases, the packing density of the peptide molecules in the coacervate increases^78,79^. As a result, the highly hydrophobic peptides are prone to producing denser coacervates, occupying less volume. This may well explain the low rim thickness for the C-terminus in general. This rationale also accounts for the relatively low *R_d_* for the N-terminus at pH 4, a condition at which it is most hydrophobic. Upon establishing the N-terminus as more hydrophilic and the C-terminus as more hydrophobic, it follows that the tick-GRP77 exhibits mixed characteristics, which we observe consistently in our results.

Building on these new insights, we revisited our earlier observations concerning the protein-depleted inverted phase inside the rim, proposing a possible link to the hydrophilicity/hydrophobicity of the peptides. Among the three IDPs, the more hydrophilic N-terminus exhibited rich rim dynamics. At pH 4, the rim thickness remained constant (*R_d_* ≈ 40 µm) with time, with no protein-depleted inverted phase observed (**Figure 4a**). However, at pH 7 and 10, the *R_d_* increased dramatically, from initial *R_d_* ≈ 20 µm to a final value of ≈ 100 µm as the phase boundary moved inwards (**Figures 4b-c**). These changes of *R_d_* over time for the N-terminus at different pH values are quantitatively shown in **Figure 4d**. At pH 4, the N-terminus is relatively more hydrophobic and hence, the spreading of rim on the glass slide was likely negligible. However, at higher pH values, the coacervate microdroplets forming in the vicinity of the phase boundary quickly fused together to form bigger droplets and finally merged with the rim (**Figure 4b-c**). Moreover, we observed prominent inverted phase both in the rim as well as within the large coacervate droplets joining the rim. Since the N-terminus becomes more and more hydrophilic with increasing pH, it is plausible that the protein-rich rim preferentially wets the hydrophilic glass slide and starts to spread, occasionally engulfing the dilute phase, resulting in the formation of the inverted phase.

**Figure 4:**
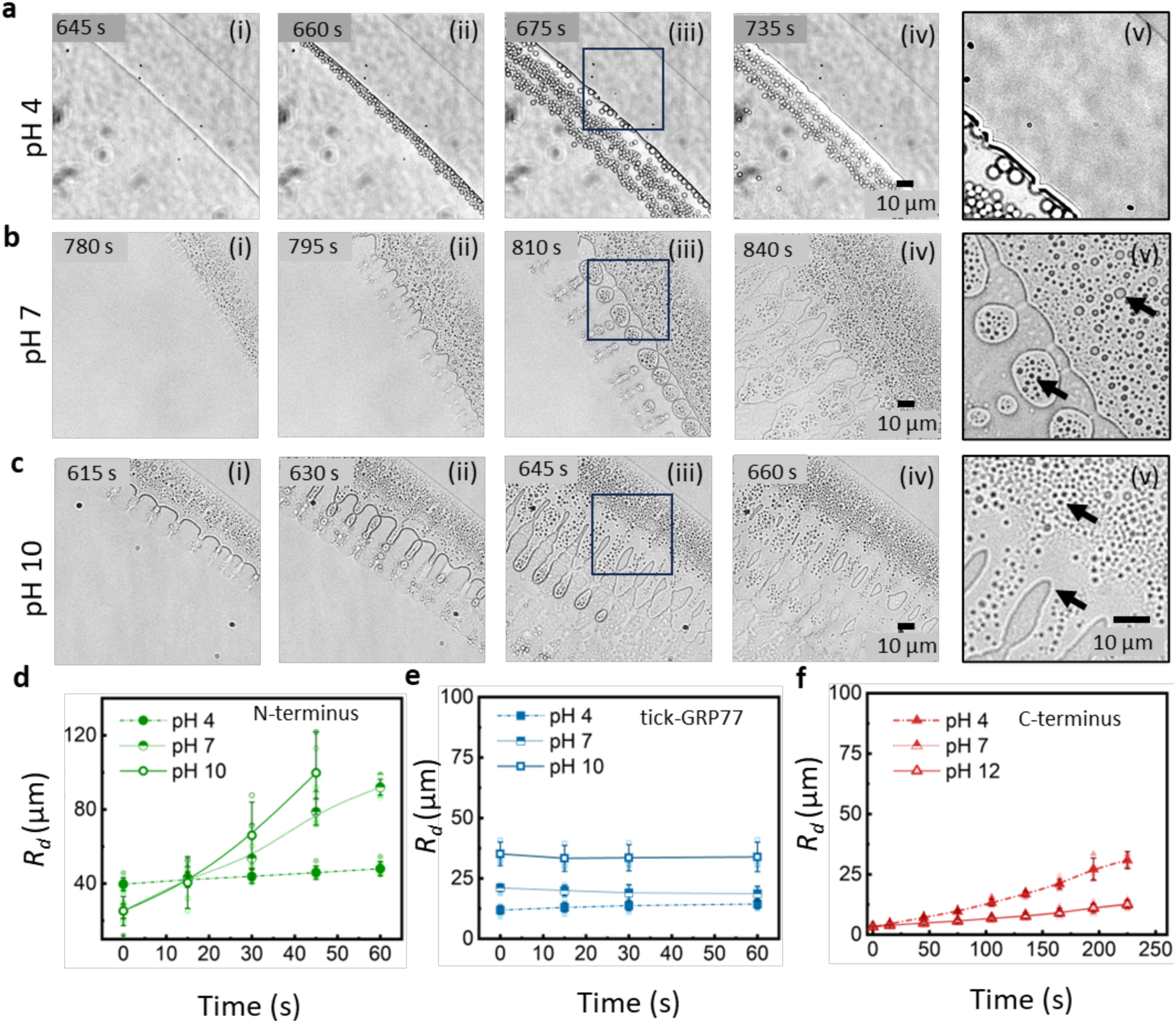
Inverted phase formation is triggered by the pH-dependent hydrophilic nature of the IDPs. **(a)-(c)** Bright-field, time-lapse images of the droplet evaporation assay for 50 µM N-terminus solutions at pH 4 (**a**), pH 7 (**b**), and pH 10 (**c**). The rim thickness in case of the N-terminus at pH 4 remains nearly constant compared to the increasing pattern for pH 7 and pH 10 with the appearance of inverted phases. **a**(**v**)-**c**(**v**) show the zoomed versions of the marked regions in **a**(**iii**)-**c**(**iii**) and the arrows indicate the inverted phases. The change in rim thickness over time for different pH values is shown for the N-terminus (**d**), tick-GRP77 (**e**) and the C-terminus (**f**). The N-terminus shows rapid rim thickness growth than the rest. The plots in **d**-**f** are represented as mean ± s.d., along with individual data points (n > 3 independent experiments in each case).

The evolution of *R_d_* for tick-GRP77 and the C-terminus are plotted in **Figures 4e-f** (time-lapse shown in **Supplementary Figures 4 and 5** ). For tick-GRP77, the *R_d_* remained constant irrespective of pH, whereas, for the C-terminus, it increased in the range of 10-25 µm. Nonetheless, in both cases, it remained below 30 µm, much lower as compared to the N-terminus. These lower *R_d_* values further showcase the relatively higher hydrophobicity of the C-terminus and tick-GRP77. Also, in case of tick-GRP77 at pH 7 and 10 (see **Supplementary Videos 2 and 3**), the protein-rich rim forms first and only at a later stage, the protein- depleted inverted phases are generated. This is in contrast with the N-terminus, where we observed the formation of inverted phase from the very beginning of rim formation. Thus we see the variation of rim patterns, R_d_, and t_i_ depend entirely on the the pH-resposive material properties of the peptides.

### The hydrophobicity of peptides determines the condensate viscosity and their interfacial activity

Next, we sought to deduce the material properties of the formed condensates in response to pH variation. We began with analyzing droplet fusion events, tracking the decay of the aspect ratio, and calculating the corresponding relaxation times, 𝜏 (see Methods for details). This simple but effective analysis has been utilized extensively to estimate the inverse capillary velocity ratio^5,80^. Reportedly,^81^ *τ* depends on three aspects, i.e., viscosity (η), interfacial energy (γ) and condensate size (d) following the relationship 𝜏 ≈ ^𝜂𝑑^.

Although all condensates underwent coalescence upon physical contact, the relaxation times varied significantly. Example curves of the decay of the aspect ratio with time for the three IDPs at pH 7 are shown in **Figure 5a**, while similar sample curves at pH 4 and pH ≥ 10 can be seen in **Supplementary Figure 6**. The curves were fitted with an exponential decay function to obtain the corresponding 𝜏 values for tick-GRP77, C-terminus and N-terminus as shown in **Figure 5b**. The 𝜏 of the C-terminus-based condensates was significantly higher than the rest, nearly 2.5- and 3-fold higher than condensates formed by tick-GRP77 and N-terminus respectively, at pH 7.

**Figure 5:**
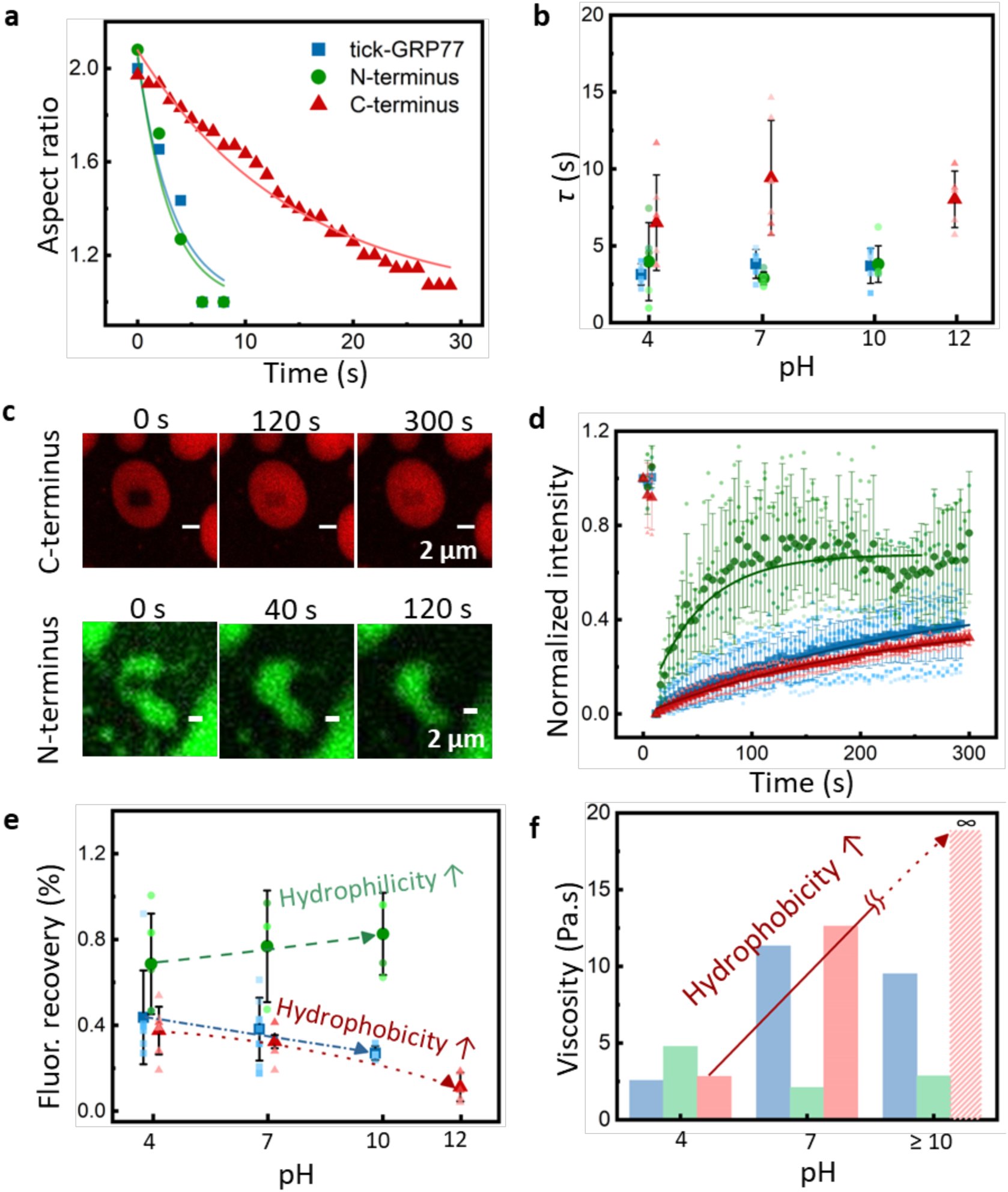
Relative hydrophobicity of the IDPs determines the material properties of the condensates. (**a**) A plot showing typical examples for the change in the aspect ratio of two fusing condensates, for each of the three IDPs at pH 7. The data was fitted using an exponential decay function (solid lines, R^2^≥ 0.9) yielding the relaxation time (τ). (**b**) Comparative bar graph showing the obtained τ values for the three IDPs at different pH values (n ≥ 5 fusion events in each case). C-terminus condensates exhibited significantly higher τ than the rest irrespective of the pH. (**c**) Representative false-colored images showing the FRAP time-lapse for the C- and N-terminus condensates at pH 4. (**d**) FRAP curves at pH 7 showing that N-terminus condensates show significantly faster recovery than the C-terminus and tick-GRP77 condensates. The solid lines show exponential fits to the datasets (R^2^ ≥ 0.8). (**e**) Plot showing the extent of fluorescence recovery for the three IDPs in response to pH. Overall higher and increasing values in response to pH for the N-terminus condensates suggest high molecular diffusivity due to their increasingly hydrophilic nature, while an opposite trend is seen in case of the C-terminus condensates. (**f**) The estimated average viscosity for the three IDPs as a function of pH. The viscosity of C-terminus condensates at pH 12 could not be determined because the recovery values were too low to be fitted and hence, a highly viscous sample is indicated. In case of FRAP experiments, 100 µM peptide concentration was used, including 5 mol% of fluorescently labelled OG488-GRP77. The plots in **b, d,** and **e** are represented as mean ± s.d., along with individual data points (n ≥ 3 independent experiments in each case). In **b** and **e**, for improved visualization, the pH values are shifted by 0.2 pH units for the three IDPs.

To further reveal the viscosity and molecular mobility inside the formed condensates, we performed fluorescence recovery after photobleaching (FRAP) experiments (see Methods for details). Higher (100 µM) peptide concentration was used for these experiments in order to have better condensate yield and thus facilitate the experiments. **Figure 5c** shows typical time-lapse images of fluorescence recovery of C- terminus and N-terminus condensates at pH 4. It appeared that the N-terminus condensates recovered significantly faster than the C-terminus condensates. The average fluorescence recovery time is plotted in **Figure 5d** for all three peptides at pH 7, where the observed difference in the recovery between N- terminus (77%) and C-terminus (32%) as well as full-length peptide (38%) is even clearer. The fluorescence recovery plots for the three IDPs at pH 4 and pH ≥ 10 are shown in **Supplementary Figure 7**.

We have shown the compilation of the fluorescence recovery of all the peptides with varying pH in **Figure 5e**. The fluorescence recovery for the N-terminus increased with increasing pH, whereas it showed a decreasing trend for the C-terminus. tick-GRP77 showed fluorescence recovery close to C-terminus and showed a decreasing trend with pH increase. We further calculated the average viscosity of condensates from the fluorescence recovery (**Figure 5f**, see Methods for details). The viscosity for C-terminus droplets at pH 12 could not be estimated as hardly any recovery was seen in order to fit the curve. Nonetheless, this clearly hints at a very high viscosity of the sample, as indicated by the patterned bar in **Figure 5f**.

These observations can again be explained based on the relative hydrophobicity of the peptides. If the peptide is highly hydrophobic, it will increase intermolecular interactions resulting in enhanced structural compactness within the condensate and a rise in its viscosity. A similar observation was reported previously where a higher KCl concentration was shown to reduce the packing compactness of complex coacervates and further reduces its viscosity^81^. Subsequently, fluorescence recovery for comparatively viscous condensates will be reduced owing to the slow diffusivity of molecules. Thus, we can infer that as the hydrophobicity of C-terminus increases with increasing pH, the formed condensates become more viscous, manifested as slower fluorescence recovery. In contrast, the more hydrophilic N-terminus forms condensates with less macromolecular compactness, which is consistent with the lower values of the estimated viscosity obtained from FRAP experiments.

Thus, multiple independent measurements revealed that tick-GRP77 and its two termini exhibited clear pH-dependent variations in their hydrophilicity/hydrophobicity. To explore the possible amphiphilic characteristics of these three peptides in response to pH variation, we prepared oil-in-water emulsions by adding octane in aqueous solution of 2 µM OG488-GRP77. **Figure 6a** shows the confocal image of the formed emulsion. While the aqueous solution shows dim fluorescence, a brightly fluorescent octane-water interface clearly shows that the tick-GRP77 preferentially adsorbed at the oil-water interface, indicating its ability to function as an interfacially active protein. To systematically study this further for the three IDPs as a function of pH variation, we conducted interfacial tension measurements using pendant drop tensiometry (see Methods for details). With the bare interfacial tension of octane-water interface measured typically at 50.8 mN/m^82^, **Figures 6b-d** display the time-dependent reduction in the base value by the three peptides (at 10 µM concentration) across the chosen pH range. tick-GRP77 reduced the interfacial tension down to 24.9 mN/m at pH 4 and further down to 20.4 mN/m at pH 10. In case of the N-terminus, the reduction was less prominent, and ranged between 32.4–39.1 mN/m. However, the C-terminus exhibited highest reduction, ranging between 25.4–13.9 mN/m. Thus, once again, we observe a completely opposite trend as a function of pH for the two termini. For the N-terminus, the interfacial tension showed the smallest reduction, indicating the lowest interfacial activity. In contrast, the C-terminus exhibited the largest reduction in interfacial tension, making it the most surface active. Not only that, for the N-terminus, the lowest interfacial tension was obtained at an acidic pH, while for tick-GRP77 and C-terminus, the lowest interfacial tension was obtained at a basic pH (see the arrows in **Figure 6c-d**). From earlier experiments, we already concluded that the N-terminus is most hydrophobic at pH 4, in contrast to the C-terminus, which shows highest hydrophobicity at pH ≥ 10. Thus, it is noteworthy that we obtained lowest interfacial tension at a pH where the peptides are most hydrophobic. Since all these IDPs are hydrophilic enough (hydrophobic/hydrophilic amino acid content ≈ 0.3), when the hydrophobicity increases, the amphiphilic nature of the protein increases, justifying the observed trend.

**Figure 6:**
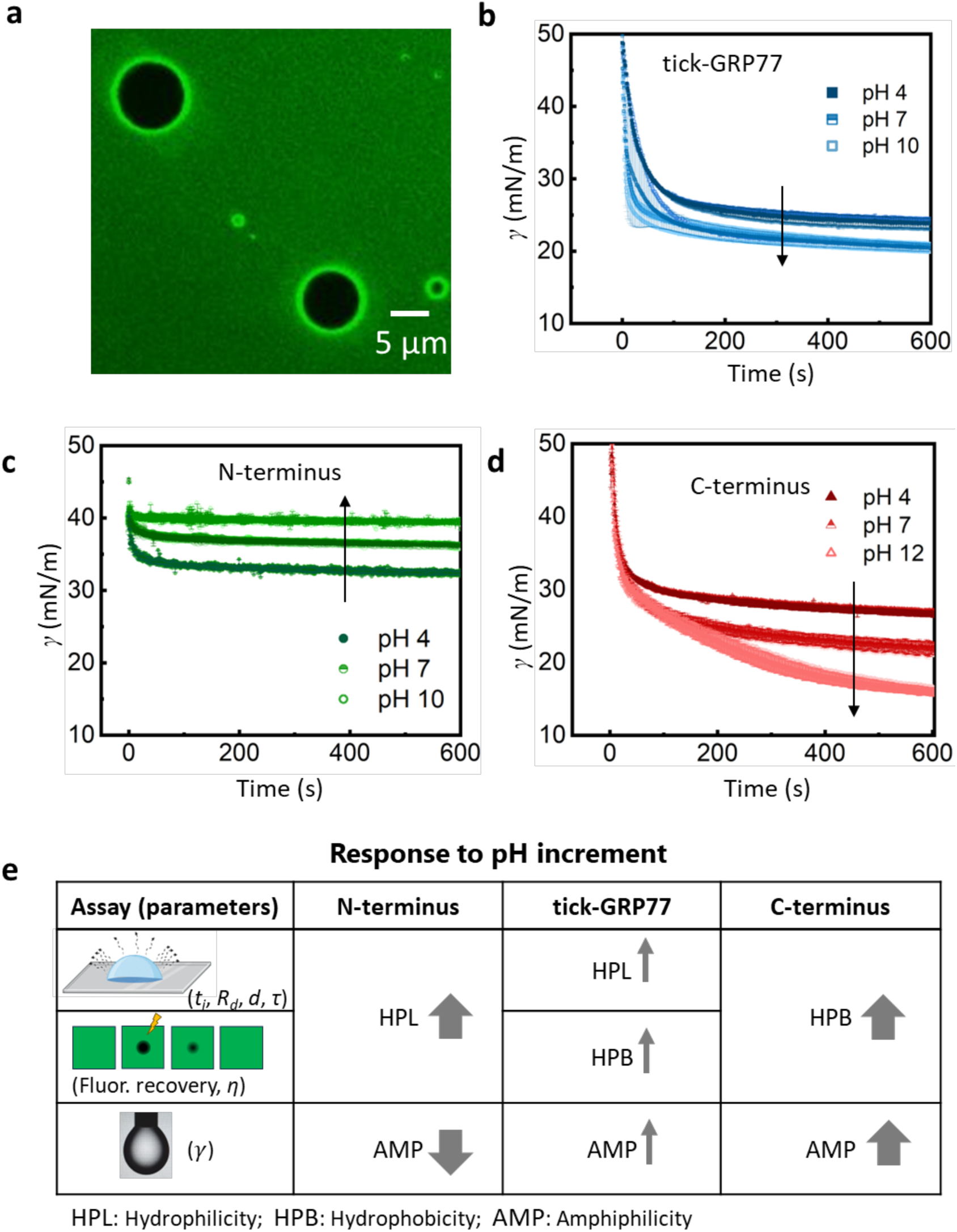
The interfacial activity of peptides is influenced by their hydrophobicity. (**a**) False-colored confocal fluorescence image of octane-in-water emulsion, showing a clear preference of fluorescently labelled OG488-GRP77 molecules to adsorb at the oil-water interface. (**b**)–(**d**) Interfacial tension measurements of the octane-water interface in presence of tick-GRP77 (**b**), N-terminus (**c**), and C-terminus (**d**) at various pH values. 10 µM peptide concentration was used for all the measurements. The plots are represented as mean ± s.d., along with individual data points (n = 3 independent experiments in each case). The arrows indicate the direction of pH increment, showing completely opposite trends for C- and N-termini. Also, the N-terminus showed the least impact on the interfacial tension, whereas the C-terminus showed the highest reduction, with tick-GRP77 showing intermediate impact. (**e**) A summary table showing the variation of material properties (hydrophilicity/ hydrophobicity/ amphiphilicity) of the three IDPs in response to pH increment as measured through droplet evaporation, FRAP, and pendent drop tensiometry assays. The thickness of the arrows indicate the strength of the trends. N- and C-termini show completely opposite traits, whereas tick-GRP77 shows intermediate character.

Although the octane-water interfacial tension in presence of the peptides is different from the interfacial tension between the condensate and the dilute phase, a comparison can still be drawn, since in both cases, the peptides decorate the interface and contribute to the interfacial properties. With these results, we can further explain the relatively higher fusion time for C-terminus condensates. Assuming similar trends as seen in **Figure 6d**, we can infer that the interfacial tension for C-terminus condensates, especially at pH 12 will be lowest. In comparison, the interfacial tension of more hydrophilic N-terminus condensates will be much higher and will increase with pH. Also, from **Figure 5f**, we see that the viscosity of C-terminus is much higher than that of the N-terminus and these values increase with pH. As fusion time is proportional to the viscosity and inversely proportional to the interfacial tension (𝜏 ≈ ^𝜂𝑑^), the high condensate fusion time of C-terminus is well justified.

Overall, we used three different experimental setups – droplet evaporation assay, FRAP, and pendent drop tensiometry to obtain interconnected results, which are compiled in **Figure 6e**. The arrows represent the change in the hydrophobic/ hydrophilic/ amphiphilic character of the IDPs as a function of pH increment and the thickness of the arrows depicts how prominent these traits were. C-terminus shows a decreasing trend for *t_i_* and *R_d_* and increasing trend for *τ* indicating its increasing hydrophobicity with increasing pH. On the contrary, N-terminus shows an opposite trend due to its increasing hydrophilicity with increasing pH. However, tick-GRP77 shows the inverted phase at more basic pH and lower *τ* similar to the N-terminus, but negligible variation in *t_i_* and *R_d_*. FRAP and pendent drop tensiometry experiments also reveal increasing hydrophobicity and amphiphilicity for the C-terminus with increasing pH and completely opposite trend for the N-terminus. Again, we see the intermediate behaviour of tick-GRP77, albeit with the values and trends closer to those of the C-terminus.

## Discussion

In this work, we demonstrate that the phase separation behavior of IDPs can be strongly dependent on the pH, by taking a model case of a glycine-rich protein (tick-GRP77) that was recently shown to exhibit phase transitions^40^. We investigated the pH-based manifestation of LLPS dynamics as well as the material properties of the formed condensates utilizing (confocal) microscopy, droplet evaporation assays, FRAP measurements, and pendant drop tensiometry. Making use of the two termini of the protein, both capable of undergoing LLPS, we found that the two halves behave antagonistic to each other as a function of pH and the roots of this behavior lie in the biochemistry of ionic and aromatic residues present within the sequence, which in turn determines their degree of hydrophobicity. The N-terminus is abundant in ionic residues but with the cations involved in cation-π interactions, anions are the primary charge contributors, making it increasingly hydrophilic as the pH increases. In contrast, being rich in cationic and aromatic residues, cations provide the key electronic contribution, making C-terminus progressively more hydrophobic with pH increase. This leads to the formaton of denser and more viscous condensates and we observe direct manifestation of this in the experiments. In addition, C-terminus exhibits the most amphiphilic character, demonstrated by the ability to reduce the oil-water interfacial tension. The N-terminus, being relatively more hydrophilic, delays condensation and produces less compact and less viscous condensates. With increase in the solution pH, its hydrophilicity increases further, which manifests in the form of an inverted protein-depleted phase. Also, N-terminus showed least reduction in oil-water interfacial tension. Interestingly, while tick-GRP77 exhibits intermediate characteristics of the two termini, it appears more hydrophobic and behaves like the C-terminus at higher pH.

This study also holds interest from the perspective of the natural occurance of the protein in tick saliva. Ticks are blood-feeding ectoparasites, and upon biting the host, they secrete a protein-rich saliva that ultimately forms a solid bioadhesive termed the ‘cement cone’ allowing them to attach to the host skin for days. GRPs are found abundantly in tick saliva and in the cement cone, as recently shown in the case of tick-GRP77^83^, and they have long been hypothesized to play a key role in cement cone formation^84^. Given the stark contrast in the pH values of relatively basic tick saliva compared to host skin and blood, it is likely that tick salivary GRPs are exposed to significant pH changes, influencing their behavior, making it important to obtain more insights about their LLPS behaviour in response to pH variation. Thus, tick-GRP77 and other tick salivary and cement cone GRPs present an excellent opportunity to study an LLPS-prone IDP that is naturally subjected to biologically relevant fundamental physicochemical triggers.

The augmentation or prevention of phase separation of these GRPs with pH variation can assist in developing tick control measures^85^. Apart from this, our work can prove useful for several biomedical applications such as tissue sealants^86,87^ as pH variation can induce the LLPS-mediated liquid-to-gel transition of the IDPs and enhance their adhesive properties. Additionally, the condensate stability can be destroyed by pH-responsiveness of the peptide releasing its contents and this can further be exploited for developing selective and targeted drug delivery^88,89^. Lastly, the pH-dependant programmable amphiphilic character of these peptides can be beneficial for stabilizing emulsions, especially in food industry^90^.

## Acknowledgements

We thank Stan van der Beelen and Leendert van den Bos from EnzyTag BV for providing tick-GRP77 and C-terminus, and Dennis Suylen from Maastricht University for helping with the synthesis of N-terminus. We also thank Xuefeng Shen for the interfacial tension measurements and Jasper van der Gucht and Soumabrata Majumdar for fruitful scientific discussions.

## Author contributions

M.N, K.A.G., and S.D. conceived the idea and designed the experiments. M.N., K.A.G., and H.I. performed the experiments. M.N. and S.D. performed data analysis. I.D. synthesized the proteins. M.N. and S.D. wrote the initial draft. M.N., K.A.G., H.I. I.D. and S.D. reviewed and edited the manuscript. I.D. and S.D. performed project administration and supervision. All authors have read and agreed to the final version of the manuscript.

## Methods

### Materials

Chemicals – Phosphate buffered saline powder, octane, sodium hydroxide, and hydrogen chloride were purchased from Sigma-Aldrich. Coverslips #1 24 mm x 60 mm were purchased from Corning®. All peptides were synthesized via solid-phase peptide synthesis (SPPS), using Boc-based SPPS and Fmoc-based SPPS. Details of the synthesis, labelling, and characterizations are given elsewhere^40^.

### Nuclear magnetic resonance spectroscopy

NMR spectra were measured on a Bruker Avance III HD 700 MHz spectrometer equipped with a three-channel TCI cryoprobe operating at a ^1^H frequency of 700.6 MHz. Sample temperatures inside the NMR probe were externally calibrated by accurately measuring the interior sample cavity temperature after manually lowering a water-filled dummy 5 mm NMR tube housing with a micro Pt100 temperature sensor, connected to a long thin cable and a datalogger positioned outside the probe.

We prepared various NMR samples of tick-GRP77 by dissolving freeze-dried powder under different buffer conditions in the pH range 3.4 to 7.4. Initial NMR spectra for structure elucidation purposes were obtained by adding MilliQ water up to a final protein concentration of 0.2 mM, with the solution slightly buffered at a low pH of 3.8, due to the presence of small amount of residual trifluoroacetic acid that was left over after SPPS.

The NMR sample of tick-GRP77 used for resonance assignment was made by dissolving 120 µg freeze-dried protein into 160 µL of MilliQ water (pH 3.8) containing 2 μL of D_2_O for deuterium lock together with ∼1 µM trace of DSS-d6 meant for internal chemical shift referencing of ^1^H and ^13^C protein chemical shifts. After mixing, the solution was transferred using a sterile plastic pipette into a standard Wilmad 3 mm NMR tube. Under these solute conditions, stable NMR spectra were obtained, leading to near complete resonance assignments made from standard 2D spectra. Standard temperature of the unbuffered low pH form of tick-GRP77 was set to 16 °C.

The sequential assignment of the tick-GRP77 protein was mostly based on DIPSI (Decoupling in the presence of scalar interactions) (80 ms mixing time), NOESY (350 ms mixing time), plus additional natural abundance heteronuclear 2D ^15^N-^1^H and ^13^C-^1^H HSQC experiments to extract carbon ^13^C and ^15^N shifts. Processing of 2D spectra was carried out with Bruker Topspin 3.6 software. Sparky 3.114 was used for analysis and manual resonance assignments of the protein.

### Microscopy and droplet evaporation assay

Stock solution of tick-GRP77, N-terminus, and C-terminus were made in MilliQ water. All the evaporation experiments were performed in phosphate buffer saline (10 mM phosphate buffer, 2.7 mM KCl, and 137 mM NaCl), with the pH adjusted to 4, 7, 10, or 12 using 1 M HCl or 1 M NaOH. A 2 µl sessile droplet of aqueous solution of protein and PBS was casted on a clean coverslip and mounted on Nikon-Ti2-Eclipse inverted fluorescence microscope and further recorded for 20-25 mins until crystallization. The coverslips were cleaned thoroguly with MQ water and dried before use. We used either Nikon Plan Apo 100x (numerical aperture, NA 1.45) oil objective or Nikon Plan Fluor 40x (NA 1.30) oil objective for all the experiments. The evolution of rim thickness was measured at regular time intervals until crystallization. The final rim thickness for all the cases was measured just before crystallization. Condensate diameters were measured using the micrographs obtained just prior to crystalization. The condensates, accumulated near the rim were considered. Using Fiji (ImageJ), a circle was drawn around the condensates, the diameter was calculated from the obtained area, and the mean droplet diameter was measured by fitting a normal distribution.

Confocal images of inverted phase and oil-in-water emulsions were recorded using Nikon C2 laser scanning confocal microscope, equipped with a 60x (NA 1.40) oil immersion objective. The samples was doped with 5 mol% OG488-GRP and was imaged using 488 nm excitation laser with excitation filter 525/50, equipped with 560 nm LP dichroic mirror and 585/65 nm emission filter.

### Droplet fusion experiments

A 2 µL drop of the protein solution under consideration was casted on a clean cover slip (24 mm × 40 mm, Corning #1.5) and was allowed to evaporate. Droplet fusion events were recorded using 40x oil objective at time interval of 1 s (exposure time of 10 ms) on Nikon Ti2 Eclipse fluorescence microscope. We used Fiji (ImageJ) to fit ellipse across the fusing condensate boundaries and estimated the aspect ratio 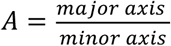. We only considered similar-sized fusing condensates to keep the analysis consistent across the samples. The change in aspect ratio as the function of time was plotted and the data was fitted using 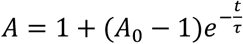; where t is the time, τ is the characteristic relaxation time, and 𝐴_0_ is the initial aspect ratio.

### Estimation of tick-GRP77 concentration within condensates using fluorescence calibration

To estimate the peptide concentration in tick-GRP77 condensates, we used confocal microscopy to calibrate the fluorescence intensity of OG488 (*I_FL_*) as a function its concentration (*C_OG488_*) and the obtained data was fitted using linear regression (*I_FL_* = 0.18 × *C_OG488_* + 0.96; R^2^ = 0.72). From the confocal images of tick-GRP77 condensates (containing 5 mol% OG488-GRP77 and using the same imaging settings as that used for calibration), we back-calculated the peptide concentration within condensates using the fit parameters. With the peptide concentration within condensates calculated at 77410.5 µM and the initial concentration at 50 µM, this estimation gave a 1548-fold increase in the peptide concentration within the condensates.

### Fluorescence recovery after Photobleaching (FRAP)

FRAP experiments were performed on Leica SP8-SMD microscope using a 63x (NA 1.2) water objective. For bleaching, the region of interest (ROI) of approximately 3 µm x 2 µm was selected inside the condensates or within the rim. The ROI was bleached using 80% laser intensity for 10 seconds and recovery of the bleached area was recorded for every 4 seconds for approximately 6 minutes. The normalized intensity of the bleached area was fitted using the equation, 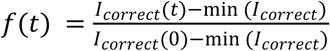, where 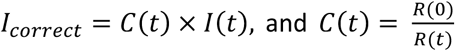. Here, *R*(*t*) and *I*(*t*) indicate the fluorescence intensity of the reference droplet at time *t* and the original fluorescence intensity of the bleached region at time *t*, respectively; min(*I_correct_*) indicates the minimum value of *I_correct_*, which is obtained right after the sample is bleached^84^. The normalized intensity was fitted using the function 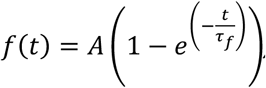, where 𝐴 and 𝜏_𝑓_ indicate the amplitude of recovery and the FRAP relaxation time, respectively. The average 𝜏_𝑓_ was estimated for each case and the average value of apparent diffusion coefficient (𝐷_𝑎𝑝𝑝_) was calculated using the formula 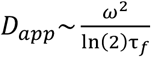 where 𝜔^2^ is the area of the bleached cross section. The average droplet viscosity, 𝜂, was estimated using 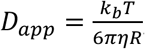, where 𝑘_𝑏_𝑇 is the thermal energy scale, and 𝑅 is the hydrodynamic radius of OG488-GRP77 (∼2.5 nm for an unfolded 7.8 kDa protein^91^).

### Interfacial tension measurements

We used a Tracker Automated Droplet Tensiometer (ADT) (Teclis, France), where a pendant droplet of 10 µL volume was generated at the tip of a SGE syringe (Trajan, Australia). The equilibrium shape of the droplet was captured by a camera, and subsequently, the data was fitted with the Young-Laplace equation using a built-in software (WINDROP_2020) to estimate the surface tension. The measurements were conducted for 600 s.

