## Supplementary Information for "pH-Responsive Phase Separation Dynamics of Intrinsically Disordered Peptides"

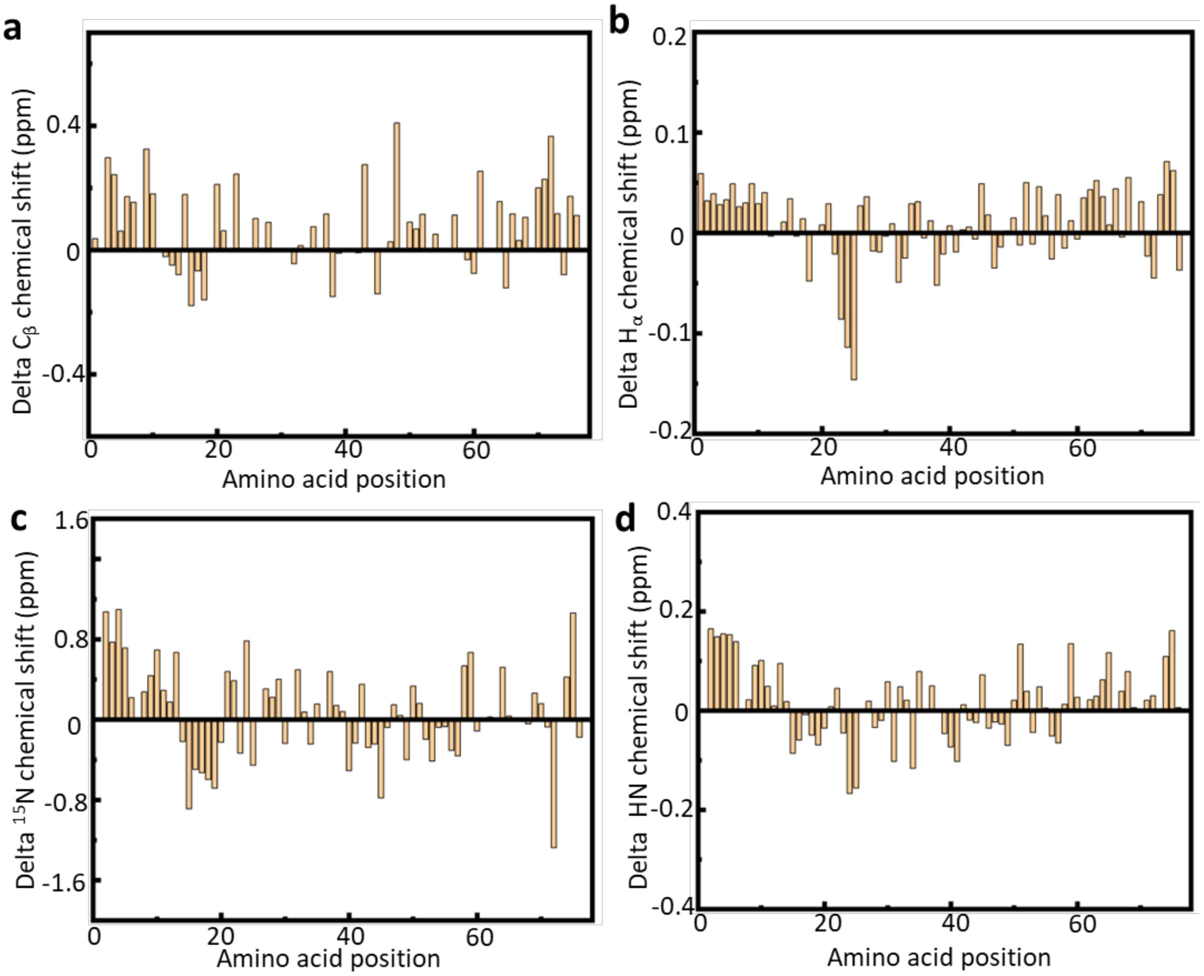


**Supplementary Figure 1. Delta chemical shifts quantifying the random coil nature of tick-GRP77.** (**a**) Delta $C_\beta$, (**b**) delta $H_\alpha$, (**c**) delta $^{15}N$, and (**d**) delta HN chemical shifts of tick-GRP77, predicting its random-coil nature. While some sequence regions display less ideal random coil compared to the $H_\alpha$ chemical shift values, these sections exhibit unusual amino acid sequence properties. For example, region GYGGPG (23-28) shows up as a negative CSI peak in the delta $H_\alpha$ profile. Also charged residues like histidine (H73) and negatively charged residues (D13, D17 and E19) show experimentally determined $^{15}N$ and $C_\beta$ chemical shifts that relatively deviate more from the predicted $C_\beta$ random coil shifts. We consider the D/E differences mainly as an artefact of the prediction method, especially when resonance positions were measured at a pH close to the isoelectric point of their charged side chains.

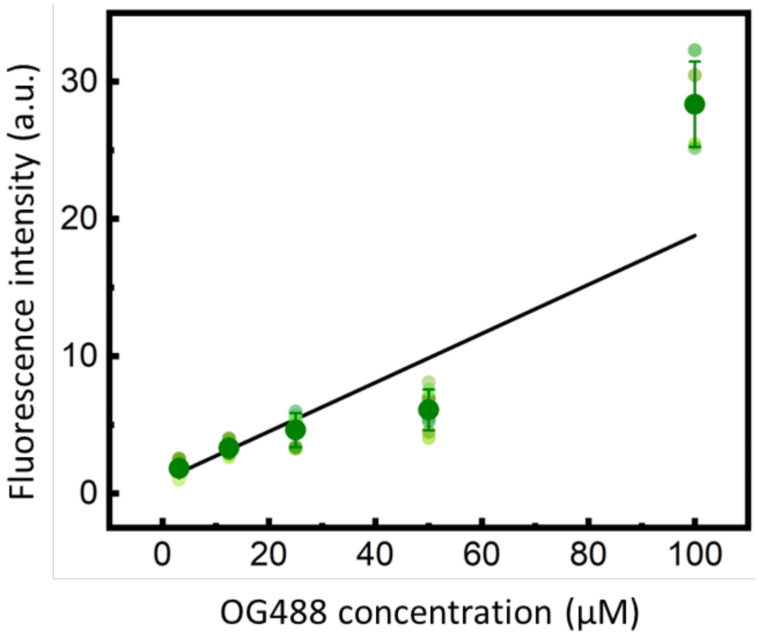


**Supplementary Figure 2. The calibration curve for estimating the tick-GRP77 concentration within condensates.** A plot of fluorescence intensity as a function of OG488-GRP77 concentration, obtained via confocal microscopy. The measurements are represented as mean ± s.d., along with individual data points ($n \geq 3$ measurements for each concentration). Solid line represents a linear fit ($R^2 = 0.7$).

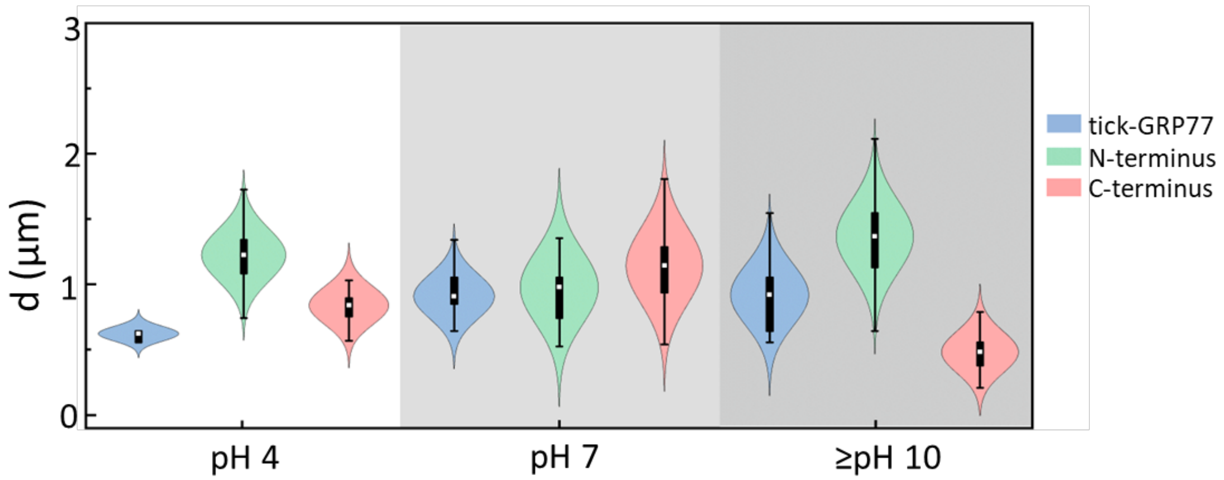


**Supplementary Figure 3. Size distribution of IDP condensates at different pH values.** The size distribution of tick-GRP77, N-terminus, and C-terminus condensates obtained during the droplet evaporation assay at different pH values (n ≥ 500 condensates in each case). The measurements are represented as violin with box plots. No clear trend or significant difference were observed. However, N-terminus condensates were generally larger than the other two, likely due to the presence of prominent inverted phase.

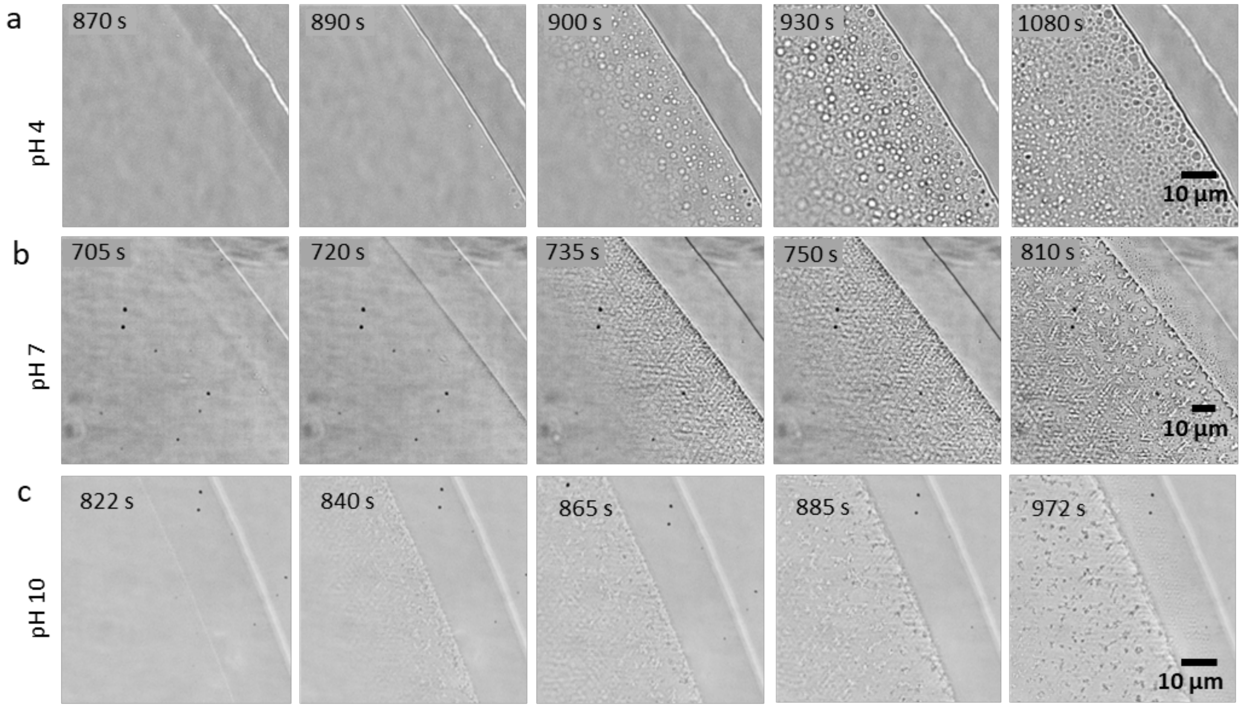


**Supplementary Figure 4**: **Rim progression for tick-GRP77 during the droplet evaporation assay.** Time-lapse images showing the boundary of an evaporating droplet of 50 µM tick-GRP77 in PBS at pH 4 (**a**), pH 7 (**b**), and pH 10 (**c**).

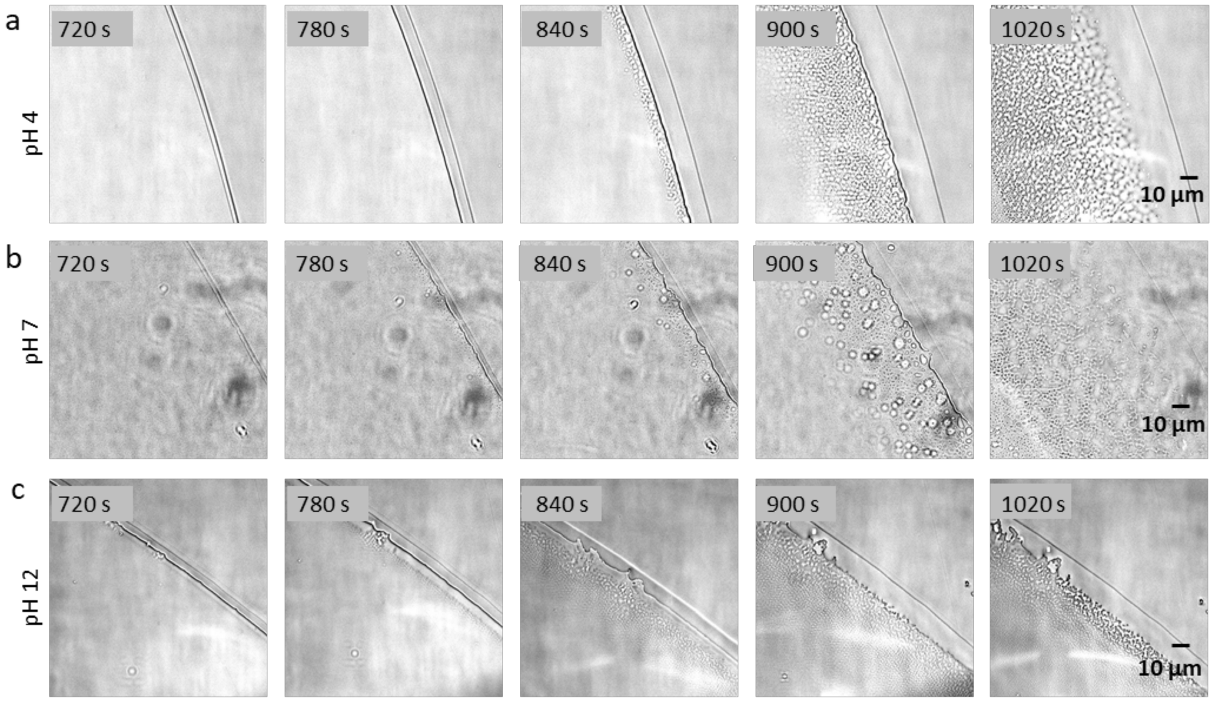


**Supplementary Figure 5**: **Rim progression for the C-terminus during the droplet evaporation assay.** Time-lapse images showing the boundary of an evaporating droplet of 50 µM C-terminus in PBS at pH 4 (**a**), pH 7 (**b**), and pH 12 (**c**).

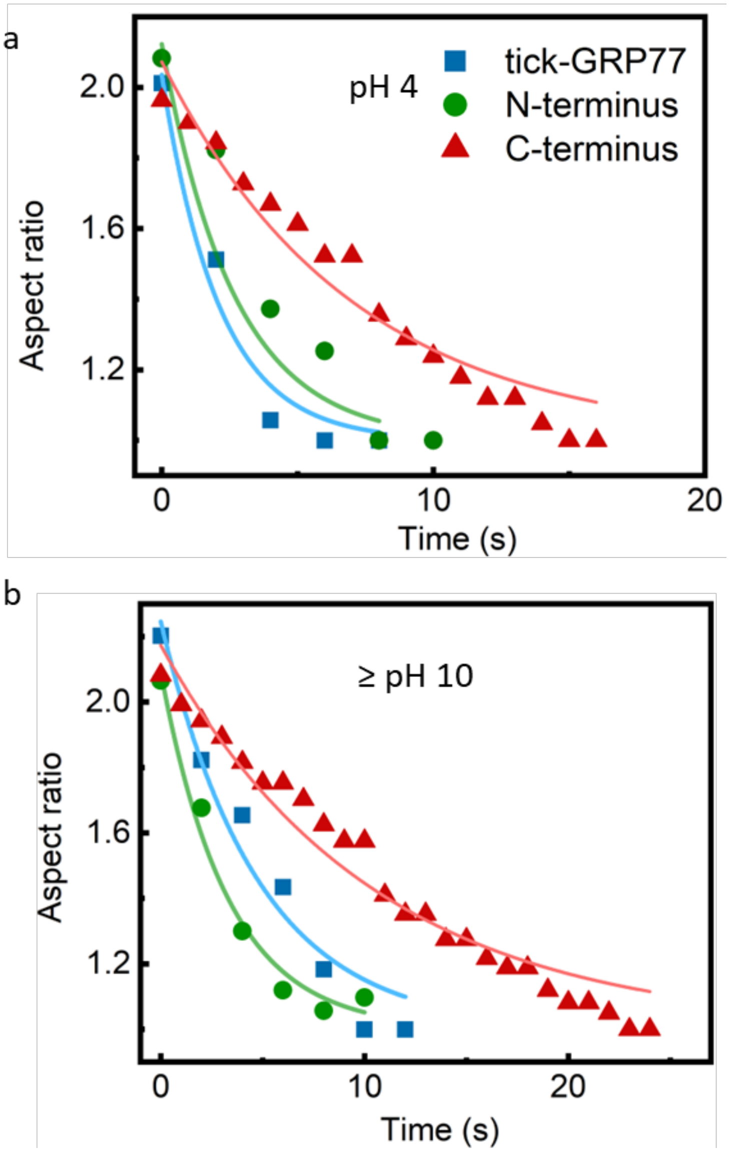


**Supplementary Figure 6. Representative droplet fusion analyses at pH 4 and 10.** Comparison of the aspect ratio decay of fusing droplets for tick-GRP77, N-terminus, and C-terminus condensates at pH 4 (**a**) and pH ≥10 (**b**). The data was fitted using an exponential decay function (solid lines, $R^2 \geq 0.9$), yielding the relaxation time. In both the cases, C-terminus condensates exhibited a significantly longer decay time, indicating slower fusion than the others.

a

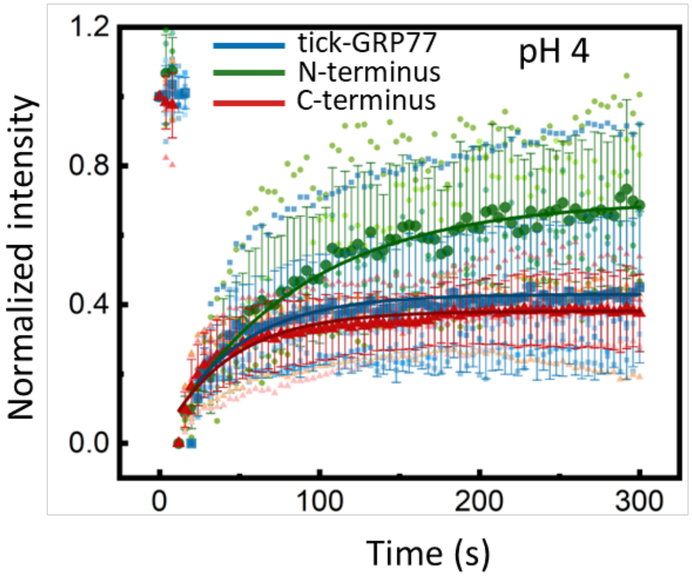


b

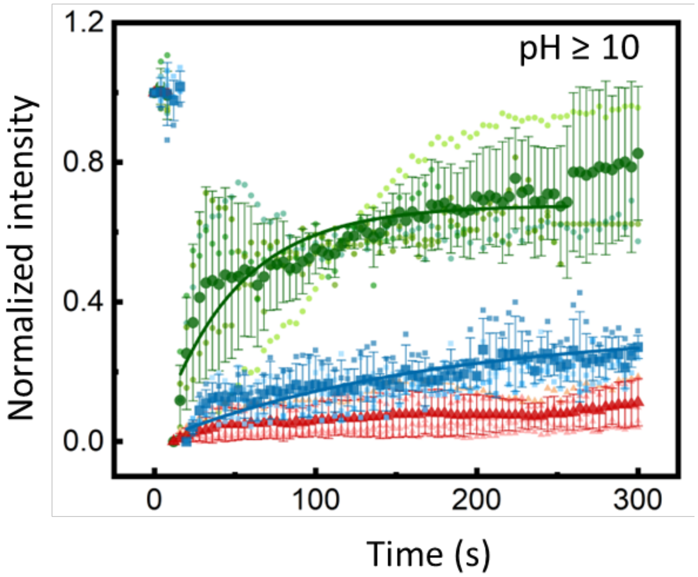


**Supplementary Figure 7: FRAP measurements of the IDP condensates at pH 4 and 10.** Fluorescence recovery curves for the three peptide condensates at pH 4 (**a**) and at pH 10 (**b**). The fitted curves are denoted by solid lines ($R^2 \geq 0.9$ for pH 4 and $R^2 \geq 0.8$ for pH ≥ 10). The N-terminus condensates showed much higher and faster fluorescence recovery at all pH values, whereas C-terminus showed the lowest recovery. The fluorescence recovery of tick-GRP77 was observed to be intermediate between the two. The plots are represented as mean ± s.d., along with individual data points (n ≥ 3 independent experiments for each condition).
